# CRISPR interference as low burden logic inverters in synthetic circuits: characterization and tuning

**DOI:** 10.1101/2020.08.03.234096

**Authors:** Massimo Bellato, Angelica Frusteri Chiacchiera, Elia Salibi, Michela Casanova, Davide De Marchi, Maria Gabriella Cusella De Angelis, Lorenzo Pasotti, Paolo Magni

## Abstract

The rational design of complex biological systems through the interconnection of single functional building blocks is hampered by many unpredictability sources; this is mainly due to the tangled context-dependency behavior of those parts once placed into an intrinsically complex living system. Among others, the finite amount of translational resources in prokaryotic cells leads to load effects in heterologous protein expression. As a result, hidden interactions among protein synthesis rates arise, leading to unexpected and counterintuitive behaviors. To face this issue in rational design of synthetic circuits in bacterial cells, CRISPR interference is here evaluated as genetic logic inverters with low translational resource usage, compared with traditional transcriptional regulators. This system has been studied and characterized in several circuit configurations. Each module composing the circuit architecture has been optimized in order to meet the desired specifications, and its reduced metabolic load has been eventually demonstrated via in-vivo assays.

## I. INTRODUCTION

The predictable composition of synthetic circuits is key to exploit the full potential of synthetic biology and enable the design of complex information processing tasks. One of the main factors leading to unpredictable behavior of synthetic circuits is cell burden [1]. The unnatural load caused by recombinant gene expression can lead to transcriptional and translational resources depletion, exerting important global effects on the functioning of the designed circuit [1,2]. Synthetic circuit designs aimed to reduce this load have been reported [3, 4, 5, 6], in which systems with higher protein expression were obtained. Experimental and in-silico methods have also been recently proposed to analyze cell burden in synthetic circuits [5, 7, 8, 9]. The use of a constitutive expression cassette for a reporter gene has been adopted as a real-time and in vivo monitor of cell capacity, to indirectly quantify cellular resource limitation [5]. This methodology was demonstrated to be more sensitive than growth rate measurement for burden quantification. Other works used the same approach to study cell burden via modelling frameworks based on electronic engineering [9] and microeconomics [8]. Different mathematical models have been proposed for the analysis of protein expression in a limited resources context [7, 10]. Such recently proposed models have been useful to identify the expression systems behaviors occurring when resources are limiting and cannot be trivially explained via simple Hill function-based activation/repression models [11]. However, such burden models still have shortcomings, e.g., they are unable to explain the possible separation of cellular resource pools among chromosome and plasmids [8] and the relationship between cell growth rate and resource pools is still lacking in such models, although recent works showed that it could have an important effect on expression dynamics [12]. While most of the literature analyzed cell burden in non-interacting gene expression systems, in-silico studies indicated that burden can largely affect the quantitative function of interconnected circuits, in which non-trivial activation and repression functions may emerge, depending on the resource usage of the individual network components [7]. This effect, not explained by traditional Hill function models, has also been confirmed in recent in vivo experiments involving simple model systems including gene regulatory networks [13,27]. An interaction graph-based theoretical framework was proposed to describe the effective interactions occurring among network modules, and eventually guide the design of circuits with different topologies [13]. The works mentioned above demonstrate the need of proper tools for cell load description and mitigation to enable a more predictable design process of synthetic circuits.

Modifications of the original CRISPR/Cas system have been proposed in the last few years to engineer new transcriptional regulators. The dead-Cas9 (dCas9) engineered protein for the targeted repression of genes, lacking nuclease catalytic activity due to single nucleotide mutations in its two cleaving domains, has been developed [14,15]. This system, known as CRISPR interference - CRISPRi, exploits targeting via single guide RNAs (gRNAs) to achieve gene silencing by steric occupancy to transcriptional elongation or binding of RNA polymerases to DNA.

Further, CRISPR/dCas9 were also fused to transcription activators, that ultimately lead to an increase in gene expression [16] despite this system so far have been developed mainly or eukaryotic cells. The Cas9 protein was also used to engineer novel fusion proteins able to act as transcriptional activators [16] or chromatin remodeling proteins [17]. Moreover, sequence-specificity of these systems implies potent selection methods of specific gene sequences in a large pool cells with different genomes, to be applied ultimately to complex bacterial ecosystems such as the human microbiome [18, 19]. Another application of this system, in which studies were performed in vivo, involves the use of a viral vector containing a combination of guide RNAs targeting retroviral LTRs and structural genes to achieve the efficient excision of pro-viral DNA from infected host cells [20], providing a viable method for the cure of diseases that until this day only have no remedy, such as AIDS and herpes. These systems rely on the multiplexing of gRNA to target several sequences while maintaining expression of the same single Cas protein; desired repression of multiple target genes with transcription factors requires complex systems and many cellular resources rendering it impractical. A recently developed CRISPR imaging system fuses GFP proteins to dCas9 for the visualization of several loci in the genome; this system, called CRISPR-Tag, allows an in-depth look at the effect of spatial-temporal organization of genes [21].

The CRISPRi system has already been adopted for the construction of logic gates as parts of interconnected synthetic circuits in different species and degree of complexity. A major advantage of CRISPRi modules, compared with transcriptional regulator proteins, is the easy programmability of gRNAs to repress the expression of any gene of interest, given the presence of an adjacent PAM motif required for system function. Here, we expand the characterization of CRISPRi repressors as logic inverter modules for synthetic circuits in *Escherichia coli* by testing their cell load mitigation in different ad-hoc constructed model systems. In fact, sgRNAs do not undergo the translation process, which is known to be the main source of cell load, thus the net load of the circuit will be only given by dCas9 and output protein expression. The designed model systems include different target promoters, copy numbers, dCas9 amounts and mutagenized gRNAs for the tuning of the NOT gate modules over a wide range of repression.

## II. RESULTS AND DISCUSSION

### A. Optimization of dCas9 expression level

For the interference system to be functional, a proper dCas9 expression level needs to be found, avoiding detrimental effects to cell growth, that is, the expression should not have a high translational demand and the synthesized protein itself should not be toxic to the cell. A popular expression plasmid (pdCas9, Addgene #44249) includes an anhydrotetracycline (aTc)-inducible dCas9. However, the steep response of the inducible promoter upon aTc addition practically resulted in a weakly reliable system for the fine tuning of dCas9 [22, 25]. As a consequence, the expression-dependent toxicity of dCas9 is still unclear since no standardized units have been used yet to characterize the effects of the expression strength. We used an HSL-inducible dCas9 expression cassette, including the P_lux_ promoter (BBa_R0062) and a strong constitutive expression cassette for the LuxR regulator, in which the P_lux_ promoter was modified to remove the last three nucleotides, which are placed downstream of the transcription start site (TSS, see section B). This system is able to achieve a strong transcription rate of the DNA downstream the P_lux_ TSS. The described HSL-inducible system was assembled downstream of a constitutive GFP expression cassette; the latter was used to monitor cell load, quantified as a drop of green fluorescence as described in [23].

To investigate the optimal expression level for dCas9, three constructs were built by combining two different constitutive promoters - with graded strengths but identical TSS (i.e. giving rise to identical transcripts) - downstream of the dCas9 expression cassette described above; those promoters drove the expression of a guide, called gPlac, targeting the P_LlacO1_ (BBa_B0011) promoter. The latter was in turn assembled upstream an RFP reporter gene, driving its expression on a co-transformed high copy plasmid (Figure 1, panel A). The choice of the constitutive promoters with different strengths (such that BBa_J23116<BBa_J23100 [26]) allowed to study the effect of increasing guide expression for several levels of dCas9 in the cell, which was tuned by varying the concentration of HSL in the medium.

**Figure-1.**
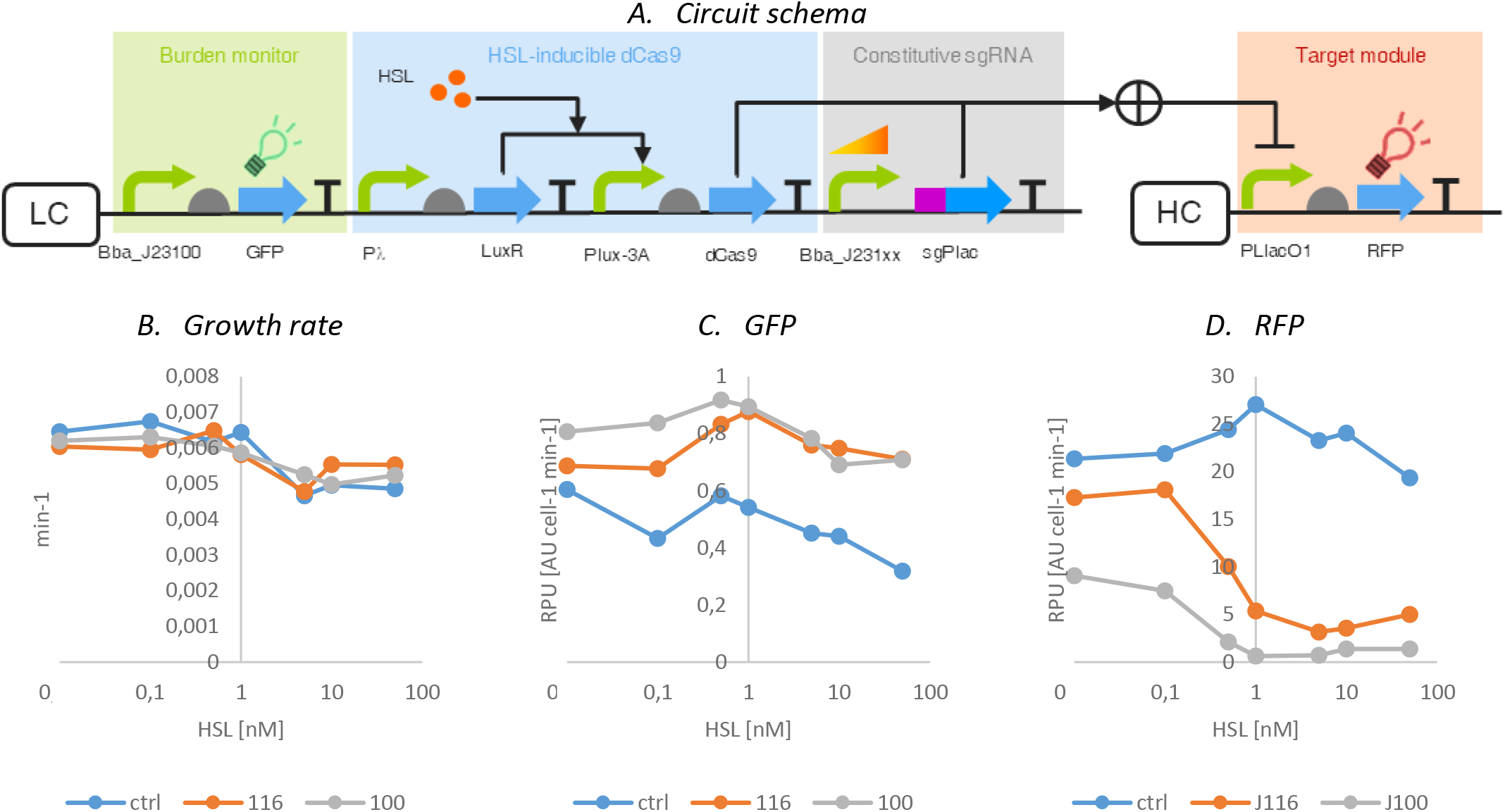
Characterization of dCas9 toxicity and repression efficiency. Circuit schema (A), growth rate (B), GFP (C) and RFP (D) signals of a system bearing a burden monitor, an HSL-inducible dCas9 expression cassette and constitutive sgRNA driven by a constitutive promoter: xx stands for 16/00 which are the BioBrick codes of the promoters used in this study (weak and strong respectively).

As a negative control, a system bearing a sgRNA - namely gPtet - designed to repress the P_LtetO1_ (BBa_B0040) promoter driven by the BBa_J23100 promoter was used; due to the orthogonality of the two sgRNA binding sites, the response was supposed to be indicative of the maximum possible expression of RFP in the absence of a complementary sgRNA, but with the same proteins and sgRNAs amounts synthesized in the cell. Through these circuits, it was possible to monitor growth rate, GFP, and RFP expression in a single time course experiment; growth rate and GFP signal were considered as proxy of cell load, while RFP was dependent to the repression capability of the sgRNA:dCas9 complex.

Dose-response experiments and steady-state data analysis were carried out as previously described [23, 24] to obtain cell growth rate and fluorescent protein synthesis rate per cell (*S*_*cell*_, in AU min^−1^) in exponential phase, as a function of inducer concentration (Figure 1, panels B,C,D).

A significant decrease in RFP output, compared with the control strain, was observed in both strains, especially the one with strong sgRNA expression, even when the dCas9 cassette was not induced, indicating the high repression efficiency of the CRISPRi repressor complex. This is likely a result of the basic activity of the HSL-based expression cassette system driving the expression of dCas9, which could be produced at low levels that are sufficient to exert significant repression.

GFP expression exhibited a slight increase at half induction of dCas9, partially attributed to the decrease in RFP expression, which represents itself a load in addition to dCas9. In other words, the burden imposed by RFP expression was relieved by the CRISPRi complex repression, allowing increased GFP expression. However, further induction of dCas9 resulted in a decrease of GFP expression and growth rate due to the high metabolic load exerted by the dCas9 protein itself.

At maximum induction of 5nM, a complete repression of the constructs with the stronger promoter (BBa J23100) was observed, while for the weak promoter (BBa_J23116) complete repression was not attained, probably due to a lack of sgRNA.

From the data gathered in these experiments, it could be firstly concluded that the CRISPRi system was functional and efficient even for low expression levels of dCas9. It is worth noting that a range of inductions was found in which dCas9 expression level showed a great efficiency without excessively affecting GFP signal and growth rate (~1 nM HSL).

For the final configuration of the circuit, a new dCas9 expression cassette was conceived by replacing the HSL-inducible system upstream the dCas9 gene with the constitutive BBa_J23116 promoter and placing the obtained device in medium copy vector (pSB3K3) in which the expression strength is comparable with the HSL-driven device with a 1-nM HSL concentration (data not shown). The combination of BBa_J23116 and medium copy plasmid to express the dCas9 gene (namely J116dCas) was expected to lead to a sufficient dCas9 level to fully repress a promoter in a high copy plasmid, provided that a proper amount of sgRNA is present in the cell, without negatively affecting growth rate or overloading the cell.

Recent literature [24] demonstrated that a high-level expression of dCas9 causes abnormal morphological changes in *E. coli*, supposed to turn from rod-shaped to filamentous in case of excessive intracellular dCas9 concentration. To verify the absence of such effects, we compared the microscope images of a strain bearing the constitutive dCas9 expression cassette described above and a control strain consisting in the same host without any exogenous plasmids (Figure 2).

**Figure-2.**
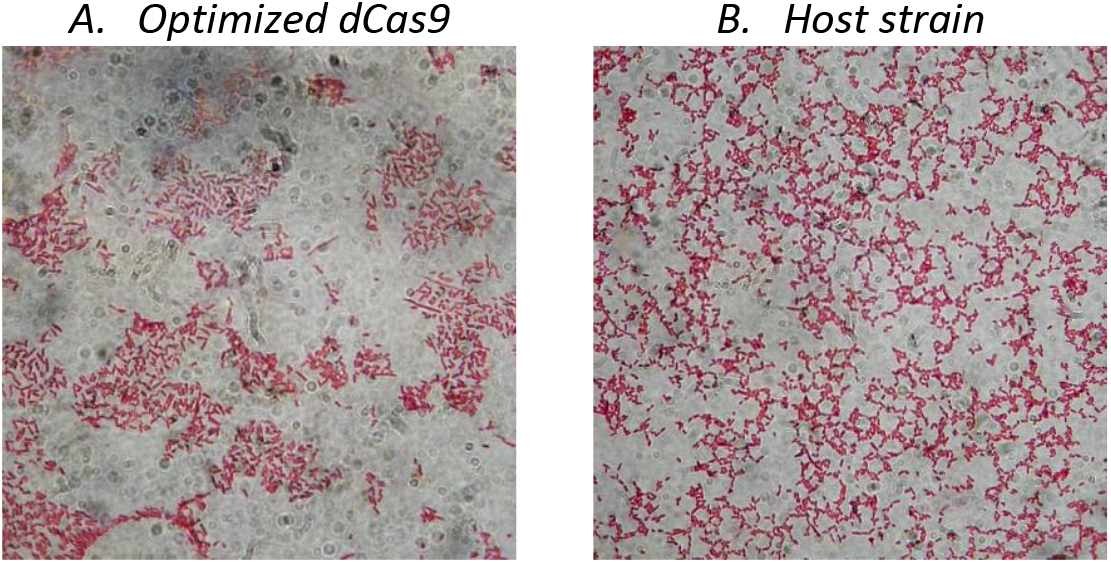
Microscopic Images of E. coli Strains with different levels of dCas9. Images were taken using the Leica bright field microscope using the 100x/1.25 oil immersion objective. Strain constitutively expressing dCas9 with an optimized expression (A), and a control strain without any exogenous plasmid (B).

Microscope images did not show any change in morphology of the cells: the construct expressing dCas9 through an optimized constitutive expression cassette showed very few filamentous cells surrounded by rod-shaped bacteria, resulting in a phenotype almost identical to the control, thus showing that a non-toxic synthesis rate was reached.

The sequence of the new dCas9 constitutive cassette was submitted to the Registry as BBa_J107202.

### B. Design of expression systems for sgRNAs

The CRISPRi complex is guided by the sgRNA molecule, whose binding with the target DNA depends on their sequence complementarity; hence, in the perspective of using inducible systems for tuning sgRNA expression, the presence of nucleotides downstream of the promoter TSS driving the expression of the sgRNA (i.e., TSS> +1) can result in elongation and possible insertions of mismatching nucleotides to the desired 20 nucleotide annealing sequence of the sgRNA. Therefore, promoters driving sgRNA expression needed to be optimized to exclude possible undesired effects. Considering the full sequences of the P_lux_ and P_LlacO1_ promoters in the Registry (BBa_R0062 and BBa_R0011, respectively), the most probable start sites of both of them were identified from sequence annotations and literature [39, 40]: for P_lux_, the TSS was in the first A nucleotide of the final sequence including three adenine nucleotides, while for P_LlacO1_ it was the last A nucleotide of the annotated promoter. This meant that the sgRNA transcribed from these promoters would include mismatches at the 5’ end of the sgRNA (i.e., three extra adenines added to the 5’ end of P_lux_ and one extra adenine to compose a 21-nucleotides guide in the case of P_LlacO1_). From other investigations on the effect of mismatches on sgRNA:DNA complementarity [30–33], it was shown that the effect of additional nucleotides to the 5’ end of a sgRNA could be marginal or significant, depending on the length of sequence extension. For this reason, in this study only promoters driving the expression of sgRNAs without extra-nucleotides were considered. Therefore, the mutagenesis of promoters driving sgRNA expression was carried out, to remove any nucleotide(s) that were initially part of the promoter but included the TSS and possibly a part of the downstream sequence. The two novel inducible devices obtained (namely P_lux-3A_ promoter and P_lac-A_), specifically implemented for sgRNA transcription, were characterized using RFP reporter to evaluate possible differences with their original sequence, by expressing the same mRNA encoding a strong RBS with RFP coding sequence used as reporter gene; this for a better understanding of the effect caused by these deletions (Figure 3). The HSL-inducible cassettes (P_lux_ and P_lux-3A_) showed a similar half-induction concentration of HSL, but significantly different maximum expression of RFP. This was probably due to the three extra-nucleotides including TSS in the P_lux_ promoter that may increase translation efficiency, rather than a change in transcriptional activity of the promoter. On the other hand, the IPTG-inducible cassettes (P_LlacO1_ and P_LlacO1-A_) behaved similarly in terms of both half-induction concentration of IPTG and maximum expression.

**Figure 3.**
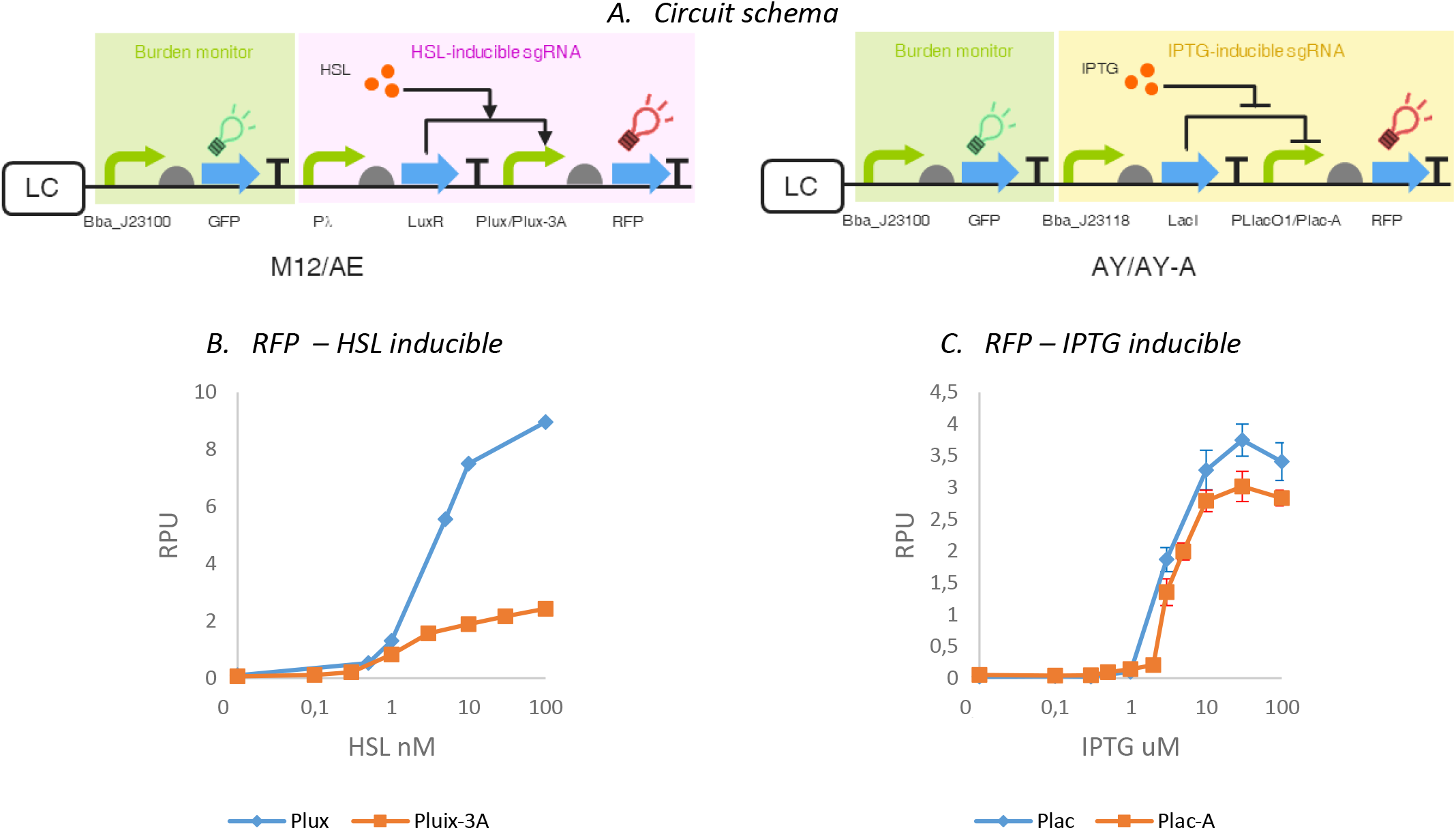
Mutagenized promoters for optimal sgRNA expression. P_lux_ and P_lux-3A_ only differ by the presence and absence of three adenine nucleotides at the end of the sequence. P_lac_ and P_lac-A_ differ only for an extra A nucleotide at the end of the sequence. Data are reported in panels B and C as mean values over at least 3 biological replicates while error bars represent the 95% confidence intervals of the mean.

Considering the overall wide tunability ranges of expression and relative induction obtained, the two systems with standardized TSS were considered suitable for sgRNA expression; however, while the modified _Plux-3A_ promoter (submitted to the Registry as BBa_J107262) was used to express the guides (see below), P_LlacO1_ was used in the downstream experiments due to the minor change in sgRNA expression (only one additional nucleotide).

### C. Characterization of a logic inverter obtained via constitutive dCas9 and inducible sgRNA

After the development of the modules illustrated above, a general standardized CRISPRi-based repression device architecture was designed and characterized by targeting P_LLacO1_. The implemented system included the following modules (Figure 4A):

- a constitutive GFP in low copy plasmid, acting as cell load measurement device, as illustrated above;
- an inducible sgRNA expression cassette in low copy plasmid obtained by using the same HSL-LuxR based expression cassette described above;
- a constitutive dCas9 expression cassette on medium copy plasmid described in the previous section;
- a constitutive target promoter driving the expression of RFP in a medium or high copy plasmid that can be repressed by dCas9:sgRNA complex.

**Figure-4.**
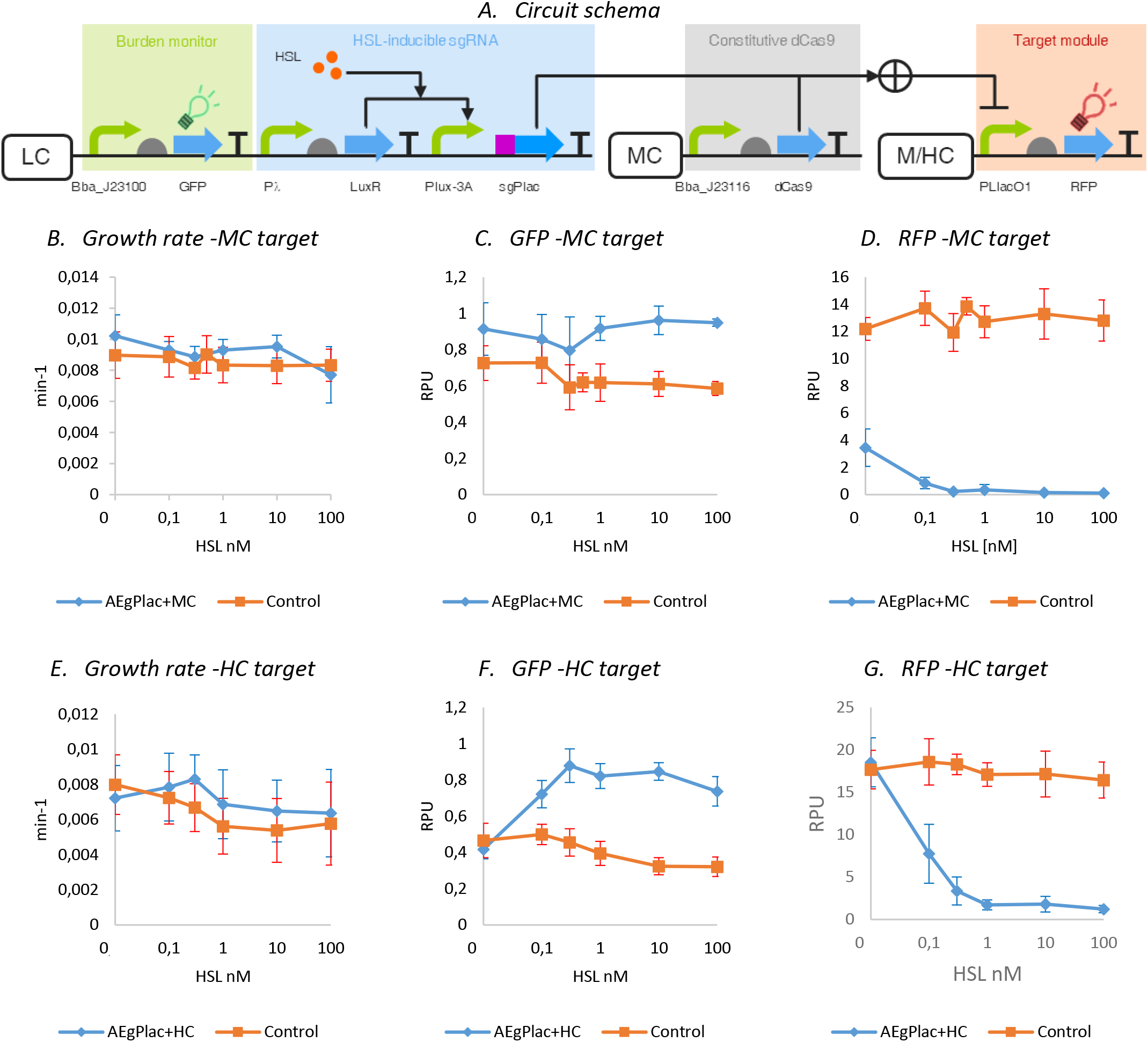
CRISPRi logic inverter with optimized dCas9 constitutive expression. The system is composed of a constitutive GFP expression cassette and an inducible sgRNA expression cassette in low copy plasmid; a constitutive dCas9 expression cassette in medium copy plasmid, and a target promoter driving the expression of RFP in medium or high copy plasmid. Control circuits bearing an sgRNA specifically targeting a different promoter under the same expression system were also considered. The measurements obtained for growth rate, GFP and RFP are shown in panels B,C,D and E,F,G for MC and HC, respectively. The x-axis represents the HSL concentration that is related to the amount of sgRNA present in the cell. Data are reported as mean values over at least 3 biological replicates while error bars represent the 95% confidence intervals of the mean.

As before, a control circuit bearing an orthogonal non-targeting sgRNA under the same sgRNA expression system was built and tested.

The optimized HSL-inducible cassette that drives the transcription of the P_LtetO1_-targeting sgRNA was therefore co-transformed with the cognate target promoter driving RFP in medium copy. Results (Figure 4, panels B,C,D) showed that the system displayed an RFP expression that was about 3-fold lower compared with the control, even without induction. Moreover, low HSL concentrations (< 1nM) were sufficient to saturate the repression, confirming the high efficiency of the CRISPRi system. Although the RFP in medium copy plasmid imposed a small burden to bacterial cells (see GFP profile for control strain), the integration of a functional CRISPRi system exhibited an improved growth rate and GFP expression compared with the control, probably due to RFP repression, which is beneficial to the host.

To improve the output range and the tunability of the system, the target was tested in a high copy vector, in an effort towards confirming and elucidating the actual repression capability of the studied systems. The system targeting the P_LlacO1_ promoter in high copy (Figure 4 panels E,F,G) in absence of HSL exhibited comparable RFP values with the control, demonstrating that the output range was successfully increased; indeed, both logic inverter strain and control displayed the same RFP value at zero induction (18RPU), which remained constant for the control and decreased readily for the functional strain, reaching the minimum value with an HSL concentration of 1nM. This result further confirmed the improved GFP expression profile of the functional system compared with the control, likely due to a higher RFP expression, as illustrated above: GFP levels were initially low for both constructs, owing to the high cell load caused by RFP expression in high copy plasmid, but it increased substantially when RFP expression was repressed by the CRISPRi complex. Growth rate showed a consistent trend in which, apart from the no-HSL data points, growth was faster for the CRISPRi-repressed strain, despite data were characterized by high variability.

### D. Transfer function tunability study

In [29], Nielsen et al. built and characterized a library of promoters of the same strength along with a library of orthogonal guides targeting specific promoters, and the inter-connectivity between parts was studied; here a complementary study in which a library of promoters of different strengths is evaluated against a single sgRNA sequence was carried out. In this context, the aim is to characterize the repression dependence on target promoter strength by using the same sgRNA sequence and target binding site, which is shared among the promoter library. A unique sgRNA (gPluxH) able to target different members of a previously constructed promoter library [27], including variable −35 and −10 sequences but identical core region, was designed. The core region of this promoter library includes the *lux box*, which is bound by the HSL:LuxR complex, thereby repressing the promoters. The gPluxH guide (Table 1) was designed to anneal a promoter sequence including the −35 boxes of the promoter library, leading to their strong repression, and was expressed through an IPTG-inducible cassette (P_LlacO1_ promoter) (Figure 5, panel A).

**Table 1:**
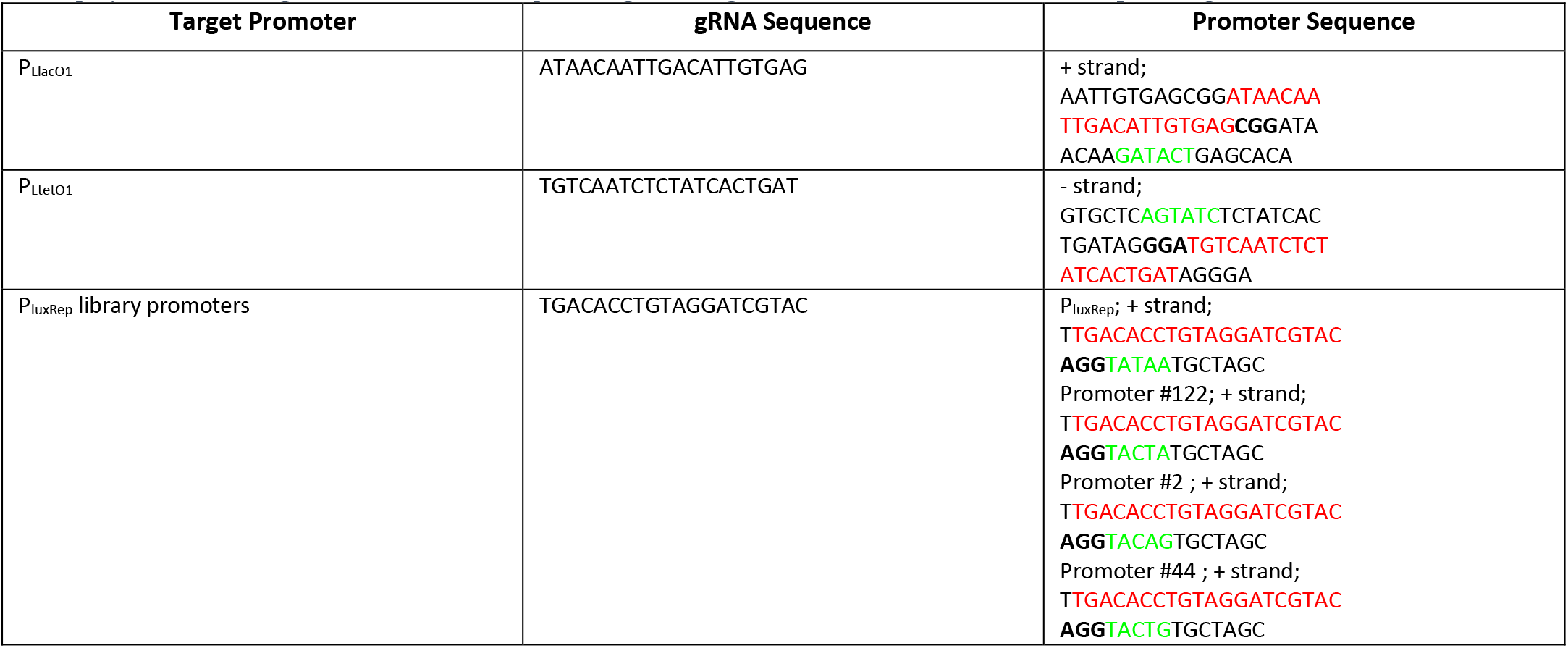
List of gRNAs and relative targets. The sequences of gRNAs conceived for the repression of target promoters are listed, as well as their respective binding locations on the target promoter. All gRNAs were designed to bind a part of the −35 box of the promoter to strongly inhibit polymerase binding. Nucleotides in red represent gRNA target, red −35Box, black bold PAM sequence, green −10Box.

**Figure 5.**
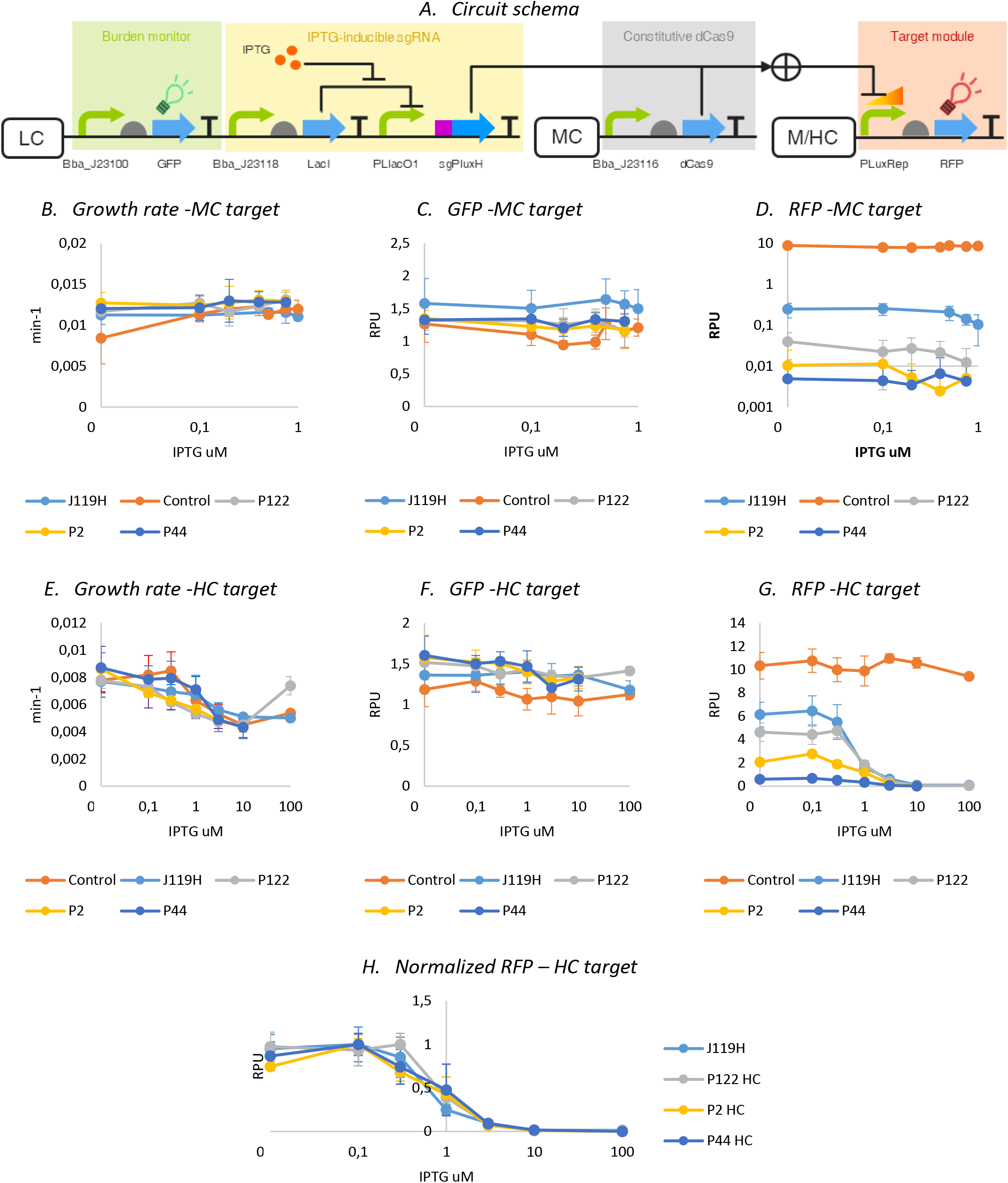
Repression of the P_luxRep_ promoter library. Target promoters had different −10 boxes, resulting in different strengths (i.e., P_luxRep_>P122>P2>P44). Measurements of growth rate, GFP and RFP expression of the transformed strains. Lines in turquoise, grey, yellow and blue represent, respectively P_luxRep_, P122, P2, P44 members of the library. The x-axis represents the concentration of IPTG in micromolar and was related to the sgRNA level in the cell. The control strain has an sgRNA targeting P_LtetO1_ that was absent in the circuit. Data are reported as mean values over at least 3 biological replicates while error bars represent the 95% confidence intervals of the mean.

A set of three promoters were chosen from the whole library presented in [27] according to their transcriptional strengths, in particular:

- BBa_J107100 (named P_luxRep_) is the strongest library member.
- BBa_J107111 (named P122) is a medium-to-high strength promoter
- BBa_J107101 (named P2) is a weak-to-medium strength promoter.
- BBa_J107105 (named P44) is a weak promoter.

Their sequences differed only at the level of the −10 box, allowing the study of the repression by a common sgRNA that strongly targets promoters with different strengths.

When the target promoters were in medium copy plasmids (Figure 5, panels B,C), all constructs maintained a stable growth rate and GFP expression over the range of the IPTG concentrations tested. RFP expression (Figure 5, panel D) was minimal for all the promoters. However, considering a logarithmic y-axis scale, it was possible to notice that the level of RFP expressed was almost conserved in terms of the ranking of promoter strengths reported in [26].

To increase the range of RFP expression and to better study the system with different configurations, the medium copy vector carrying the target promoters was replaced with a high copy vector, leading to a higher expression level of RFP (Figure 5, panels E,F,G).

As expected, the high copy targets showed increased RFP values, comparable with the medium copy ones in terms of ranking, and this increase in RFP was accompanied by a decrease in growth rate. In particular, the promoters showed ranking consistent with the original publication previously illustrating their strength, with the J119H promoter being the strongest one. At high IPTG concentrations, RFP becomes strongly repressed. This means that gPluxH in the tested condition can exert a high repression strength and range as a function of IPTG (related to the amount of sgRNA transcribed), leading to well-tunable RFP levels. Nonetheless, RFP expression from the circuit with the strong P_luxRep_ promoter showed slightly lower level than the control (which includes the same setup, but with a non-specific sgRNA), meaning that the basic activity of P_LlacO1_ in the off-state (i.e., without IPTG) was sufficient to exert a relevant repression. The respective descending order of promoter strengths were measured to have RFP values of 6.1 RPUs, 4.6 RPUs, 2.1 RPUs, and 0.6 RPUs, respectively.

To evaluate the transcriptional strength-dependent sgRNA-mediated repression among the tested constructs, their RFP expression were normalized to the maximum RPU reached by each curve in the no-IPTG experimental condition (Figure 5, panel H). The normalized curves showed no relevant change in repression function shape among the four promoters, thereby suggesting that transcriptional strength does not play a role in sgRNA efficiency in terms of percent activity repression. On the other hand, as expected, absolute RFP expression depends on promoter strength in both non-repressed and repressed state. Specifically, the normalized curves showed similar IPTG concentration leading to half-maximum repression and slope.

Growth rate and GFP expression did not show relevant IPTG-dependent and promoter strength-dependent variation, except for the control strain, in which GFP is lower than the other tested constructs, consistent with the highest RFP expression shown by the control due to its non-repressed condition.

Taken together, the results suggested that the characterization of sgRNAs in one context can be used to quantify their repression strength, which can be generalized to other target promoters with the same sgRNA binding region. Despite results showed that no percent repression difference can be seen, they again showed that repression strength is highly dependent from the specific sgRNA: the ones designed to repress P_luxRep_ showed higher repression strength than the one targeting P_LlacO1_. This might be a combined effect of sgRNA:dCas9 dissociation constant with target DNA, and the different promoter systems that have been used to drive sgRNA expression.

### E. Conclusions

In this work, different investigations focused on the common goal of facilitating the use of CRISPRi to build novel customized gene regulatory networks with low cell load were provided.

Specifically, the first investigation aimed at finding a low-toxicity high-efficiency trade-off for the constitutive expression of dCas9, to obtain an optimized fully functional minimal burden dCas9 expression cassette. The results showed that a low transcription rate on a medium copy plasmid, which is co-transformed in engineered bacteria in all the quantitative experiments done in this work, was successful to efficiently implement a logic inverter module based on sgRNA. After the optimization of dCas9 expression, a number of synthetic circuits in which sgRNAs were expressed by inducible promoters to target RFP expression by binding to its upstream promoter were built and characterized. The tests carried-out with target promoters in medium- or high- copy plasmids, showed to be all functional in terms of repression. As expected, the target copy number plays a role in the output curves by showing tighter repression by sgRNAs when target is in medium -copy compared to high -copy, in which the tunability range of repression was higher. The CRISPRi system is more exploitable in the choice of target than other specific protein transcription factors. It is worth noting that, despite all the tested sgRNAs designed in this work act as repressors, the repression efficiency strongly depends on the sequence, e.g., the sgRNA designed to target the library of P_luxRep_ promoters exerts a much stronger repression than the one targeting P_LlacO1_. In some cases, repression was so strong that the output showed negligible RFP expression for any sgRNA level. This means that the basic activity of inducible promoters driving sgRNAs was sufficient to produce sgRNA levels with good repression capabilities, and it might also depend on the promoter system used to drive the expression of sgRNAs. GFP data, used as a proxy of cell load, showed that no relevant burden affects the cells upon sgRNA expression. Importantly, metabolic load was much lower than the one observed in some repression systems based on widely used protein repressors (e.g., TetR), thereby suggesting that the use of CRISPRi in synthetic circuits as a low-burden alternative to transcription factors can be successful.

Lastly, to better understand the possibly occurring interplay between CRISPRi system and commonly found genetic circuit designs, we studied the effects on sgRNA repression capability as a function of transcriptional activity of target promoter. It resulted that the fold-repression caused by sgRNA was not dependent from promoter strength: an sgRNA targeting a small library of promoters with diverse activity but sharing the same core sequence (in which the guide RNA binds) showed similar repression curves. This result enables the predictable re-use of previously designed sgRNAs to target different promoters sharing a conserved sequence. In general, despite no relevant cell burden was detected in the sgRNA-based circuits, some of the systems showed slight GFP and growth rate drops which were not expected. Similar effects are still under study by many groups and their full understanding is expected to improve the predictability of bottom-up designed biological systems. Having seen that the constitutive expression of dCas9 was effective in this configuration, further decreasing its expression could allow for a larger range of RFP values, although the use of several sgRNAs in the same circuit may require additional dCas9 pool.

Taken together, these results suggested that CRISPRi can be successfully used to design low-burden solutions. Nonetheless, further studies are needed to fully elucidate and quantitatively characterize system behavior in all the considered genetic contexts and for additional sgRNAs case studies. Mathematical modeling investigations aimed at predicting the behavior of such genetic circuits can be of support to increase knowledge on the behavior of the devices herein developed. Noticeably, this work has offered an enrichment to the CRISPRi toolkit for *Escherichia coli*, providing new data and efficiency demonstration for logic inverters that can be used in synthetic circuits exerting low cell load. Targeting was carried out by taking into account widely used regulated promoters that are herein repressed by sgRNAs instead of their cognate repressor protein. This enables to exploit such experimental setup to add regulations and inputs to existing logic gates based on popular inducible systems. Despite investigations of CRISPRi systems in organisms other than *E. coli* falls out of the scope of this study, the results of this work will facilitate the future use of the CRISPR/dCas9 system as customizable and low-burden regulation strategy for synthetic circuits and complex metabolic pathways.

## III. METHODS

### A. Strains, reagents and cloning

A list of all the constructs used and developed in this study is reported in Table 2.

**Table 2:** Table of constructs. The table shows the BioBrickTM codes for the constructs built in the study as well as their composition.

| Name | Construct | Purpose |
| --- | --- | --- |
| dCas9 | BBa J107200 + pSB3K3 | Nuclease null Sp.Cas9 protein from Addgene plasmid #44249 |
| tracr | BBa J107201+pSB1A2 | tracr RNA with tetraloop for Sp.dCas9 from Addgene plasmid #44251 |
| A37 | BBa J23100 + BBa B0032 + BBa E0040 + BBa B0015 + pSB4C5 | GFP-based metabolic burden Monitor |
| AE dCas9 | A37 + BBa R0051 + BBa B0030 + BBa C0062 + BBa B1006 + BBa R0062 + BBa J107200 + pSB4C5 | LuxR-driven dCas9 inducible cassette for toxicity measurement |
| M12 Inv | A37 + BBa R0051 + BBa B0030 + BBa C0062 + BBa B1006 + BBa R0062 + BBa B0034 + BBa E1010 + BBa B0015 + pSB4C5 | LuxR-driven RFP inducible cassette for toxicity comparison with monitor |
| AE d116gPtet | AE dCas9 + BBa J23116 + gPtet + BBa J107201 + pSB4C5 | dCas9 efficiency characterization on tetR target promoter – low sgRNA |
| AE d100gPtet | AE dCas9 + BBa J23100 + gPtet + BBa J107201 + pSB4C5 | dCas9 efficiency characterization on tetR target promoter – medium sgRNA |
| AE d119gPtet | AE dCas9 + BBa J23119 + gPtet + BBa J107201 + pSB4C5 | dCas9 efficiency characterization on tetR target promoter – high sgRNA |
| AE d116gPlac | AE dCas9 + BBa J23116 + gPlac + BBa J107201 + pSB4C5 | dCas9 efficiency characterization on lacI target promoter – low sgRNA |
| AE d100gPlac | AE dCas9 + BBa J23100 + gPlac + BBa J107201 + pSB4C5 | dCas9 efficiency characterization on lacI target promoter – medium sgRNA |
| AE d119gPlac | AE dCas9 + BBa J23119 + gPlac + BBa J107201 + pSB4C5 | dCas9 efficiency characterization on lacI target promoter – high sgRNA |
| E62 | BBa R0040 + BBa B0034 + BBa E1010 + BBa B0015 + pSB3K3 | tetR target promoter in medium copy |
| I13521 | BBa R0040 + BBa B0034 + BBa E1010 + BBa B0015 + pSB1A2 | tetR target promoter in high copy |
| E52 | BBa R0011 + BBa B0034 + BBa E1010 + BBa B0015 + pSB3K3 | LacI target promoter in medium Copy |
| A33 | BBa R0011 + BBa B0034 + BBa E1010 + BBa B0015 + pSB1A2 | LacI target promoter in high copy |
| J116dCas | BBa J23116 + BBa J107200 + pSB3K3 | Constitutive dCas9 expression cassette in medium copy |
| E20 | BBa R0051 + BBa B0030 + BBa C0062 + BBa B1006 + BBa R0062 + BBa B0034 + BBa E1010 + BBa B0015 + pSB4C5 | Template for optimized LuxR-inducible guide expression cassette |
| AE tracr | A37 + BBa R0051 + BBa B0030 + BBa C0062 + BBa B1006 + BBa J107202 + BBa B0034 + BBa E1010 + BBa B0015 + BBa J107201 + pSB4C5 | Template for guide mutagenesis in LuxR expression cassette |
| AE-3A | A37 + BBa R0051 + BBa B0030 + BBa C0062 + BBa B1006 + BBa J107202 + BBa B0034 + BBa E1010 + BBa B0015 + pSB4C5 | Optimized LuxR-inducible expression cassette characterization |
| AYA | A37 + BBa J23118 + BBa B0034 + BBa C0012 + BBa B0015 + BBa R0011 + BBa B0034 + BBa E1010 + BBa B0015 + pSB4C5 | LacI-inducible expression cassette characterization |
| AY Atracr | A37 + BBa J23118 + BBa B0034 + BBa C0012 + BBa B0015 + BBa R0011 + BBa B0034 + BBa E1010 + BBa B0015 + BBa J107201+pSB4C5 | Template for guide mutagenesis in LacI-inducible expression cassette |
| AY-A | A37 + BBa J23118 + BBa B0034 + BBa C0012 + BBa B0015 + BBa J107203 + BBa B0034 + BBa E1010 + BBa B0015 + pSB4C5 | Mutagenized LacI-inducible guide expression cassette characterization |
| AEgPtet | A37 + BBa R0051 + BBa B0030 + BBa C0062 + BBa B1006 + BBa J107202 + gPtet + BBa J107201 + pSB4C5 | LuxR-inducible guide cassette for repression characterization |
| AEgPlac | A37 + BBa R0051 + BBa B0030 + BBa C0062 + BBa B1006 + BBa J107202 + gPlac + BBa J107201 + pSB4C5 | LuxR-inducible guide cassette for repression characterization |
| AYgPtet | A37 + BBa J23118 + BBa B0034 + BBa C0012 + BBa B0015 + BBa R0011 + gPtet + BBa J107201 + pSB4C5 | LacI-inducible guide cassette for repression characterization |
| AY-AgPtet | A37 + BBa J23118 + BBa B0034 + BBa C0012 + BBa B0015 + BBa J107202 + gPtet + BBa J107201 + pSB4C5 | Mutagenized LacI-inducible tet guide for repression characterization |
| AYgPluxH | A37 + BBa J23118 + BBa B0034 + BBa C0012 + BBa B0015 + BBa R0011 + gPluxH + BBa J107201 + pSB4C5 | LacI-inducible guide cassette for repression characterization |
| dCasE62 | BBa J23116 + BBa J107200 + BBa R0040 + BBa B0034 + BBa E1010 + BBa B0015 + Psb3k3 | Constitutive dCas9 and tet target promoter in medium copy |
| dCasE52 | BBa J23116 + BBa J107200 + BBa R0011 + BBa B0034 + BBa E1010 + BBa B0015 + pSB3K3 | Constitutive dCas9 and lac target promoter in medium copy |
| dCasJ119H | BBa J23116 + BBa J107200 + BBa J107100 + BBa B0034 + BBa E1010 + BBa B0015 + pSB3K3 | Constitutive dCas9 and PluxRep target promoter J119H or PluxRep in medium copy |
| dCasP122 | BBa J23116 + BBa J107200 + BBa J107111 + BBa B0034 + BBa E1010 + BBa B0015 + pSB3K3 | Constitutive dCas9 and PluxRep target promoter #122 in medium copy |

The *E. coli* TOP10 (Invitrogen) strain was used as a host for cloning and quantitative assays. The strain was transformed by heat shock at 42°C, according to manufacturer’s instructions. LB medium was used during plasmid propagation. Antibiotics were always added to maintain plasmids in recombinant strains: ampicillin (100mg/l), kanamycin (50mg/l) or chloramphenicol (12.5mg/l). Long-term bacterial stocks were prepared for all the engineered strains by mixing 750ul of a saturated culture with 250ul of 80% glycerol, and stored at −80°C.

All the plasmids used in this study were constructed through BioBrick Standard Assembly [34] and conventional molecular biology techniques. As a result, standard DNA junctions (TACTAG upstream of coding sequences, TACTAGAG otherwise) are present between assembled parts. The BioBrick basic or composite parts used for DNA assembly were retrieved from the MIT Registry 2008-2011 DNA Distribution [35] except for P_luxRep_, which was constructed in a previous study [26] and the new ones conceived in this work.

DNA purification kits (Macherey-Nagel), restriction enzymes and T4 DNA ligase (Roche), Phusion Hot Start II PCR kit and T4 polynucleotide kinase (Thermo Scientific) were used according to manufacturer’s instructions. Plasmids were sequenced via the BMR Genomics DNA analysis service (Padova, Italy) and Eurofins Genomics. Oligonucleotides for mutagenesis were obtained from Metabion International AG and Eurofins Genomics.

M9 supplemented medium (11.28g/l M9 salts, 1mM thiamine hydrochloride, 2mM MgSO_4_, 0.1mM CaCl_2_, 0.2% casamino acids and 0.4% glycerol) was used in quantitative experiments. HSL (#K3007, Sigma Aldrich) was dissolved in deionized water to prepare a 2mM stock, stored at −20°C.

### B. Mutagenesis

All the guides used in this work were designed via the CRISPR tool of Benchling, setting the guide length to 20 nucleotides, GCA_00005845.2 as reference genome, using the Optimized Score from Doench, Fusi et al., 2016 and keeping unchanged all the others default settings. Mutagenesis with divergent primers was adopted to customize gRNAs and to delete nucleotides after the transcription start sites of the used promoters, when indicated. For gRNA sequence insertion, tailed 40 nucleotide primers were used such that 20 nucleotides composed the gRNA sequence and the other 20 nucleotides annealed the 5’end of non-annealing part of the sgRNAs. Deletion were obtained via amplification of the desired part of the plasmid, excluding the nucleotides that needed to be deleted. The experimental protocol was as follows:

- Template plasmid DNA was purified; the Phusion Hot Start Flex II was used according to manufacturer protocol; primer pairs used in the study are listed in Table 3.
- The PCR cycle was run and followed by Dpn1 (Roche) digestion of the methylated template DNA; primer annealing temperature was calculated on the free online tool offered by New England Biolabs with parameters set as the standard for Phusion polymerase;
- PCR products were separated in a 1% agarose gel, and extracted and purified;
- Blunt-end DNA fragments were phosphorylated by Polynucleotide Kinase (PNK - New England Biolabs) and ligated by T4 ligase in the same reaction; a 20μL reaction mix was composed of:

- A maximum of 50ng of DNA brought to a volume of 17μL with deionized water;
- 2μL of ligase buffer;
- 1μL of PNK;

**Table 3:**
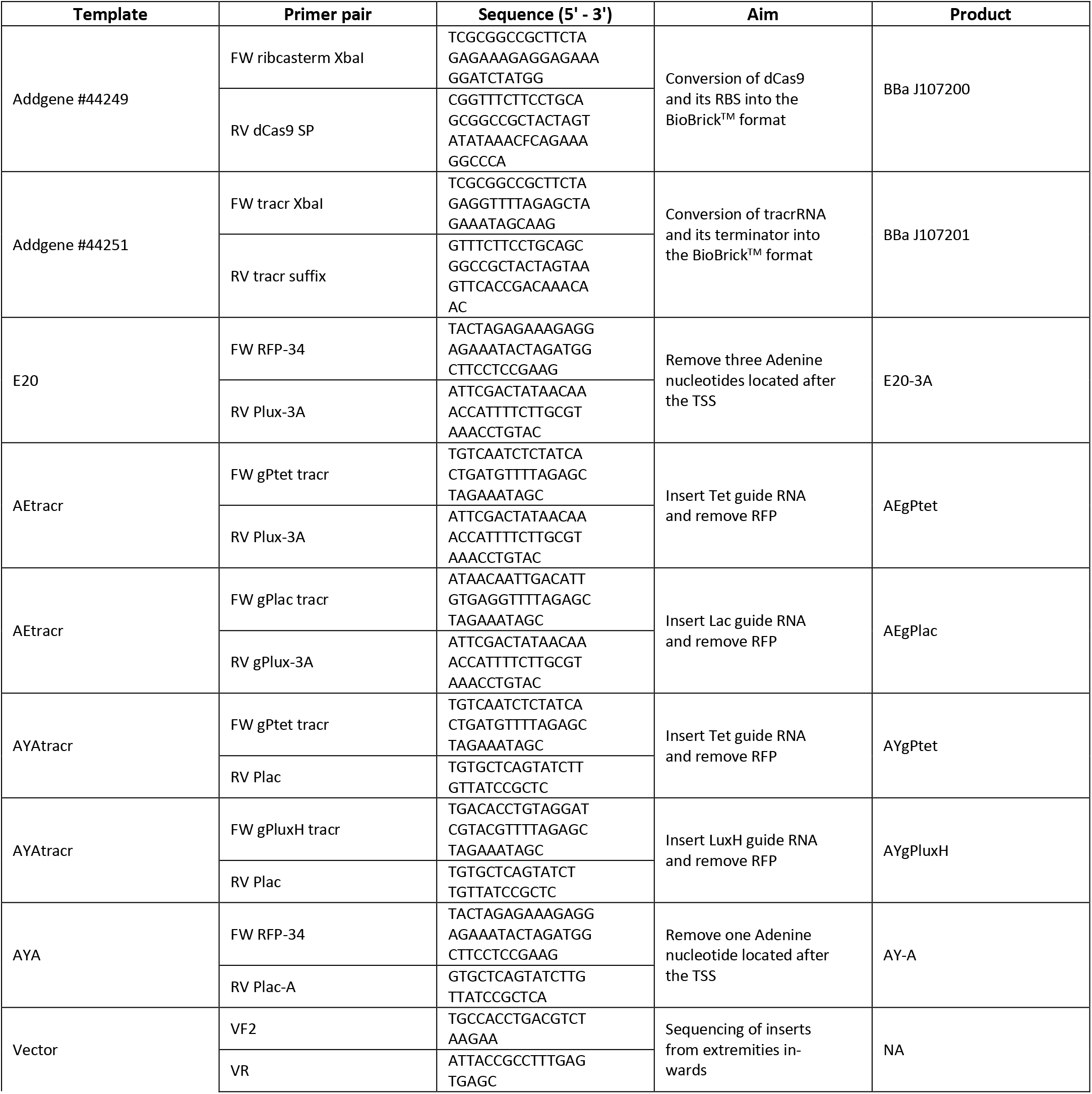
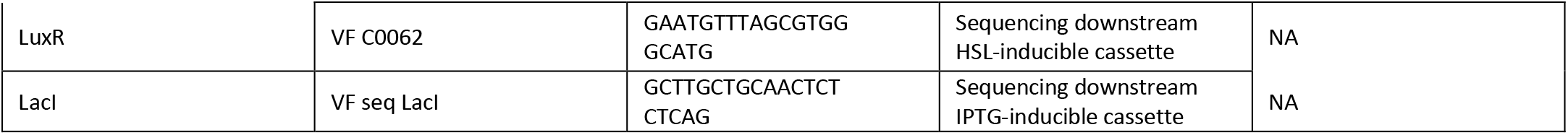
Table of primers. Primers used in the study are listed and their purpose is stated.

The reaction was allowed to proceed at 37°C for 20 minutes, then 1μL of ligase was added to the reaction mix and incubated for 16h at 16°C; finally, PNK and T4 Ligase were deactivated at 75°C for 10 minutes, and the same protocol for transformation and stock preparation described upstream was employed.

### C. Amplification and standardization of CRISPRi elements

To facilitate all the assemblies carried out in this work and to support the re-use of constructed parts in future works, the two main elements of the CRISPRi system (i.e. dCas9 and tracrRNA) were PCR amplified from Addgene plasmids #44249 and #44251 with the convergent primer pairs FW_ribcas_XbaI, RV dCas9 SP and FW_tracr_XbaI, RV_tracr_suffix, respectively, to convert them into the BioBrick format. The resulting sequences were digested and ligated in the BioBrick compliant pSB3K3 plasmid via Biobrick Assembly [36] (avoiding digestions with EcoRI of the pars bearing dCas9 gene, naturally containing this restriction site).

### D. Circuits characterization

Fluorescence and absorbance of recombinant bacteria incubated in a microplate reader were measured over time as previously described [27, 37, 38]. Briefly, bacteria from a glycerol stock were streaked on a selective LB agar plate. After 16h to 20h incubation at 37°C, 1ml of selective M9 was inoculated with a single colony. After 21h incubation at 37°C, 220rpm, in an orbital shaker, cultures were 100-fold diluted in a final volume of 200ul in a 96-well microplate. HSL (2ul) was added when required, to reach the desired final concentration.

Cultures were not placed in the external wells of the plate to avoid intensive evaporation during incubation. The microplate was incubated with lid in the Infinite F200Pro microplate reader (Tecan) and it was assayed via kinetic cycle: 15s linear shaking (3mm amplitude), 5s wait, absorbance (600nm) measurement, fluorescence measurements, 5min sampling time. RFP and GFP fluorescence were measured with a gain of 80 with the 535-620nm and 485-540nm filter pairs, respectively. Control wells were always included, as described in the following Data processing Section, to measure the background of absorbance and fluorescence, and to provide internal control references for relative activity calculations.

### E. Data processing

Data analysis and graphs were carried out via Microsoft Excel and Matlab R2017b (MathWorks, Natick, MA). Linear regression (for growth rate measurement) was carried out via the Matlab *regress* function.

Raw absorbance and red fluorescence time series were blanked by background subtraction as previously reported [6,28] to obtain OD600 and RFP time series, proportional to the per-well cell density and fluorescent protein number. Sterile medium and a non-fluorescent TOP10 culture were used as absorbance and red fluorescence background, respectively.

Since a significant cell density-dependent and growth rate-dependent autofluorescence was previously reported for GFP measurements with our experimental setup, green fluorescence was blanked via a different procedure, similar to a previous work [28]. In the previous work, each recombinant strain was implemented and tested in 2 different versions, with and without GFP expression cassette, which did not cause any relevant difference in cell load and growth rate. Here, to avoid the construction of GFP-negative strains for each condition, we estimated the cell density- and growth rate-dependent autofluorescence as 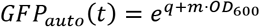, in which OD600 is the cell density of the culture, and *m* and *q* describing the linear relation between ln(OD_600_) and GFP autofluorescence level. The *m* and *q* coefficients are growth rate (μ)-dependent parameters, with a function defined as:

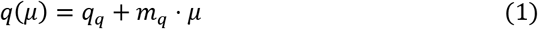

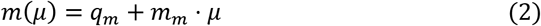

Therefore, the autofluorescence expression becomes:

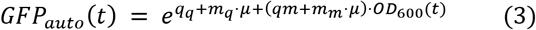

Using Equations 1–3, the four coefficients were fitted from autofluorescence data of different cultures exhibiting diverse growth rates. This expression served as a calibration curve that provides autofluorescence values (that are subtracted from raw green fluorescence values), given OD600 and μ.

### F. Microscope assay of cell wealth

The Leica DMLS Type 020-518.500 bright field microscope was used to take images of bacterial cells for morphological analysis. Bacteria were magnified with the 100x/1.25 oil immersion objective, and static pictures were taken using the Nikon COOLPIX 4500 digital camera. For sample preparation, the following protocol was employed:

- long-term bacterial stocks were streaked on selective LB agar plates;
- colonies were used to inoculate 500µL of M9 medium supplemented with the appropriate antibiotic(s)
- 20uL of culture were fixed on a glass slide by heating over a Bunsen burner;
- Fixed cells were stained for 90s with a 0.5% Safranin solution diluted in deionized water;
- The staining solution was washed away with an adequate amount of running tap water and left 10 minutes to air dry under a fume hood;
- The slide was mounted with a cover slip with 60uL of EUKITT mounting medium and left to solidify under a fume hood;

## Author Contributions

MB, LP and PM conceived of the study. MB drafted the manuscript. MB, ES, MC, AFC conducted the experiments. MB, ES, LP and PM analyzed the data PM and MGCD provided reagents and materials. All authors read and approved the final manuscript.

## Funding

This work was supported by Fondazione Cariplo through the grant 2015–0397 “Conversion of industrial bio-waste into biofuels and bioproducts through synthetic biology”. The funders had no role in study design, data collection, analysis and interpretation, decision to publish, or preparation of the manuscript.

## Abbreviations

CRISPRi: Clustered Regularly Interspaced Short Palindromic Repeats interference
GFP: Green Fluorescent Protein
HSL: N-oxohexanoyl-L-homoserine lactone
IPTG: Isopropyl-β-D-1-thiogalactopyranoside
OD600: optical density at 600 nm
PCR: polymerase chain reaction
RBS: ribosome binding site
RFP: Red Fluorescent Protein
rpm: rotation per minute

## Notes

### Competing Interest Statement

The authors have declared no competing interest.

